# Early Treatment Response in First Episode Psychosis: A 7-Tesla Magnetic Resonance Spectroscopic Study of Glutathione and Glutamate

**DOI:** 10.1101/828608

**Authors:** Kara Dempster, Peter Jeon, Michael MacKinley, Peter Williamson, Jean Théberge, Lena Palaniyappan

## Abstract

Early response to antipsychotic medications is one of the most important determinants of later symptomatic and functional outcomes in psychosis. Glutathione and glutamate have emerged as promising therapeutic targets for patients demonstrating inadequate response to dopamine-blocking antipsychotics. Nevertheless, the role of these neurochemicals in the mechanism of early antipsychotic response remains poorly understood. Using a longitudinal design and ultra-high field 7-Tesla magnetic resonance spectroscopy (MRS) protocol in 53 subjects, we report the association between dorsal anterior cingulate cortex glutamate and glutathione, with time to treatment response in drug-naïve (34.6% of the sample) or minimally medicated first episode patients with non-affective psychosis. Time to response was defined as the number of weeks required to reach a 50% reduction in the PANSS-8 scores. Higher glutathione was associated with shorter time to response (F=4.86, *P*= .017), while higher glutamate was associated with more severe functional impairment (F=5.33, *P*= .008). There were no significant differences between patients and controls on measures of glutamate or glutathione. For the first time, we have demonstrated an association between higher glutathione and favourable prognosis in FEP. We propose that interventions that increase brain glutathione levels may improve outcomes of early intervention in psychosis.

## Introduction

Early treatment response has been identified as one of the most robust predictors of longer-term clinical outcomes in schizophrenia.^1^ In particular, lack of early response appears to be strongly indicative of subsequent non-response,^2^ failure to achieve remission^3^, and higher rates of treatment discontinuation^4^. Approximately one third of patients with schizophrenia are considered to be treatment resistant^5^, with the majority of these (23-34%) failing to respond appreciably to dopamine-blocking antipsychotic medications from their first episode of psychosis (FEP).^6, 7^ Nevertheless, the neurochemical mechanism of early response is poorly understood, precluding efforts to prevent or reduce the rates of treatment failure and persistent disability.

The FEP is characterized by a relative state of glutamatergic excess.^8, 9^ Elevated anterior cingulate cortex (ACC) glutamate has been found to be inversely correlated with striatal dopamine synthesis in patients with FEP^10^. Given that the elevated striatal dopamine synthesis relates to better treatment response^11^ in psychosis, the observed glutamatergic excess has been considered to be an index of reduced treatment responsiveness in psychosis^12^.Elevated anterior cingulate cortex (ACC) glutamate has been directly associated with lack of remission in certain samples of chronic^13, 14, 15^ or first-episode schizophrenia^16, 17^ [UK sample], but this has not been a consistent observation. For example, in a sample of patients with established schizophrenia, Iwata et al (2018) ^18^ reported no difference in dorsal ACC glutamate levels between treatment-responsive and resistant groups. Similarly, the samples in 2 out of 3 sites in another study showed no glutamate excess in patients with FEP who did not achieve remission by 1 month^17^. Nevertheless, relative glutamatergic excess appears to be specific to early stages of illness^8^, and relates to more severe symptoms at presentation^17^, as well as grey matter decline^19^, cognitive^20^ and functional^16, 17^ impairments. The lack of dopamine elevation seen in some patients may explain their lack of response to dopamine blocking medications.

Glutathione (GSH), the brain’s most prominent intracellular antioxidant has been suspected to play a key protective role in free-radical-mediated damage to neurons^21^, giving rise to the redox dysregulation hypothesis of schizophrenia^22^. MRS studies have found a small but significant GSH deficit in the ACC in patients with schizophrenia^23^, indicating the presence of subgroups of patients with different redox profiles^24^. The most prominent reduction in GSH seems to occur particularly in patients with persistent residual symptoms, indicating that low levels of GSH may be associated with poor response to antipsychotics^25^. Furthermore, N-acetyl-cysteine (NAC), a precursor of GSH, appears to increase the rate of symptomatic response when used as an adjunct to antipsychotics^26^. Glutamate is a precursor of GSH while GSH acts as a neuronal reservoir for glutamate synthesis^27^. As a result, when neuro-glial metabolic integrity is intact, glutamate and GSH levels remain tightly linked in the brain. Glutamatergic excess can result in neurotoxic oxidative stress^28^, while a concomitant elevation of GSH may provide a neuroprotective ‘gate-keeping’ effect^29^, thus a strong covariance may be a marker of a healthy state. Nevertheless, repeated or prolonged exposure to excess glutamate can deplete GSH levels^30^. Furthermore, the GSH-glutamate homeostasis may also be disrupted in patients with schizophrenia due to deficiencies in GSH synthesis^31^, leading to reduced GSH-glutamate covariance in patients with FEP.

In this study, we use ultra-high field 7T MRS for the first time to test the relative contribution of ACC GSH deficiency and glutamatergic excess in predicting early treatment response in FEP. Given the gatekeeper role of GSH in tackling oxidative stress^31^, we expected GSH to be a more critical determinant of early treatment response in FEP. We hypothesized that FEP patients with higher GSH levels would demonstrate faster symptom reduction upon starting antipsychotic treatment (hypothesis 1). As not all patients with FEP will be able to increase GSH in accordance with glutamate levels, we expected a reduction in the strength of correlation between the GSH and glutamate levels in patients compared to healthy controls (hypothesis 2). Furthermore, in light of the excitotoxic theory of acute schizophrenia^32^, we expected both reduced GSH and increased glutamate levels to predict impaired Social and Occupational Functioning at the onset of illness (hypothesis 3).

## Methods

### Participants

The sample consisted of 37 new referrals to the PEPP (Prevention and Early Intervention for Psychosis Program) at London Health Sciences Centre between April, 2017 and January, 2018. All potential participants provided written, informed consent prior to participation as per approval provided by the Western University Health Sciences Research Ethics Board, London, Ontario. Inclusion criteria for study participation were as follows: individuals experiencing FEP, and having received antipsychotic treatment for less than 14 days in their lifetime. A consensus diagnosis was established using the best estimate procedure^33^ for all participants after approximately 6 months by 3 psychiatrists (KD/LP and the primary treatment provider) based on the Structured Clinical Interview for DSM-5^34^. Participants meeting criteria for bipolar disorder with psychotic features, major depressive disorder with psychotic features, or suspected drug-induced psychoses were excluded from further analyses. Antipsychotic medications were chosen by the treating psychiatrist and the patient and/or their substitute decision maker in a collaborative manner. There was no specific protocol in place regarding switching antipsychotic medications in this naturalistic sample. If an individual did switch medications, this was noted and the reasons for switching were recorded. Over the course of the follow-up period for this study, 9 individuals switched antipsychotic medications, and in all cases, the reasons for switching were related to side effects. In accordance with current national guidelines for the treatment of FEP, all individuals were offered the option of treatment with a long acting injectable at the earliest opportunity^35^.

Healthy control subjects were recruited through the use of posters advertising the opportunity to participate in a neuroimaging study involving tracking outcomes following FEP. Healthy control subjects had no personal history of mental illness, and no family history of psychotic disorders. Group matching with the FEP cohort for age, sex, and parental education was maintained. Exclusion criteria for both the FEP and healthy control groups involved meeting criteria for a substance use disorder in the past year according to DSM-5^36^ criteria (this was based on self-report for controls, and in addition clinical assessment and urine drug screening done at the point of clinical assessment in suspected cases for patients), having a history of a major head injury (leading to a significant period of unconsciousness or seizures), having a significant, uncontrolled medical illness, or having any contraindications to undergoing MRI. Clinical Measures

While the proportion of FEP patients in remission at any given time appears to be relatively consistent, it is often not the same individuals who remain in remission at each time point^37^. The use of absolute criteria in defining remission is highly dependent on initial illness severity, with individuals with a higher initial symptom burden being much less likely to achieve remission^38^. As a result, we studied the continuous measure of time to response as the primary clinical outcome of interest, and used the cross-sectional remission criterion as a secondary measure of interest.

The 8 items of the Positive and Negative Syndrome Scale capturing the core symptoms critical in defining remission (PANSS-8^39^) was administered at baseline, 2 weeks, 4 weeks, and at every clinical encounter thereafter on a 2-4 weekly basis. The PANSS-8 has acceptable internal consistency and comparable sensitivity to early improvement in psychotic symptoms^40^ relative to the PANSS-30^41^. The time to achieve a 50% PANSS-8 improvement from baseline^42^, sustained for at least 2 consecutive visits 2 weeks apart, was used as a continuous measure of treatment response. A 50% symptom improvement from baseline roughly equates to a Clinical Global Impression-Schizophrenia (CGI-S^43^) scale score of “much improved” thus, is clinically meaningful^44^. Relative PANSS8 improvement was calculated as (PANSS8_baseline_ - PANSS8_endpoint_)/(PANSS8_baseline_ - 8) in order to adjust for the minimal possible PANSS8 score^45^. All patients were observed clinically for a period of at least 6 months, and no patients failed to reach this milestone within this time frame.

We also assessed binary remission status after the first month of treatment (remission or not in remission). Symptomatic remission was allocated based on remission criteria proposed by Andreasen et al (2005)^39^ which categorize remission as achieving scores of mild (3) or less on all PANSS8 items, without any stipulation of a duration criteria, in line with Egerton et al^16, 17^. Finally, social functioning was assessed at baseline using the Social and Occupational Functioning Assessment Scale (SOFAS^46^).

### Medication Adherence

Individuals were treated with long-acting injectable (LAI) medications whenever clinically appropriate. Patients taking LAI’s received their injection from a nurse at the PEPP clinic and therefore, it was known if an individual had missed, or was late for their scheduled dose. Assessments of medication adherence were also recorded at each clinical encounter, taking into account information provided by the patient, their family, and/or case manager using a 5-point rating scale (ranging from 0 for individuals not taking medication to 4 for those being adherent 75–100% of the time). This measure has been found to correlate with pill counts^47^. We only included subjects who had >75% recorded adherence.

### ^1^H-MRS

Metabolite concentrations (glutamate and GSH) were estimated using single-voxel 1H-MRS data acquired with a Siemens/Agilent MAGNETOM 7.0T head-only MRI (Siemens, Erlangen, Germany; Agilent, Walnut Creek, California, USA) using an 8-channel transmit/32-channel receive head coil at the Centre for Functional and Metabolic Mapping of Western University in London, Ontario. A 2.0 × 2.0 × 2.0 cm (8cm^3^) ^1^H-MRS voxel was placed in the bilateral dorsal ACC (see figure 1) using a two-dimensional anatomical imaging sequence in the sagittal direction (37 slices, TR=8000ms, TE=70ms, flip-angle (α)=120°, thickness = 3.5mm, field of view = 240×191mm). The posterior end of the voxel was set to coincide with the precentral gyrus and the caudal face of the voxel coincided with the most caudal location not part of the corpus callosum. The angulation of the voxel was determined to be tangential to the corpus callosum. A total of 32 channel-combined, water-suppressed spectra were acquired using a semi-LASER ^1^H-MRS pulse sequence (TR=7500ms, TE=100ms) during each scan session, while participants were at rest and asked to stare at a white cross on a black screen for 4 minutes. Water suppression was achieved using the VAPOR preparation sequence^48^, and water-unsuppressed spectra were acquired for spectral quantification and line shape deconvolution reference. The 32 spectra were corrected for frequency and phase drifts as described in Near et al (2015)^49^ prior to averaging and lineshape deconvolution using QUECC^50^. Residual water peaks were removed from the averaged spectrum using HSVD^51^. Metabolite quantification was acquired using Barstool^52^. Water-subtracted spectra were modelled using the fitMAN, a-prior-knowledge based minimization algorithm, and a quantification template including 17 metabolite spectral signatures derived from simulation^52^. Our fitting template included 17 metabolites (alanine, aspartate, choline, creatine, GABA, glucose, glutamate, glutamine, glutathione, glycine, lactate, myo-inositol, N-acetyl aspartate, N-acetyl aspartyl glutamate, phosphorylethanolamine, scyllo-inositol, and taurine). Importantly, at this long echo time, no macromolecules were included in the spectra as their signal had decayed below noise level. Metabolite concentrations were corrected for gray and white matter volumes using the anatomical MRI images and previously described methods^53^. All spectra and spectral fit were inspected visually for quality and Cramer-Rao lower bounds (CRLB) were assessed for each metabolite. The MRS metabolite estimates were not known at the time of clinical outcome characterization. See the Supplement for further details on the MRS methods.

**Figure 1.**
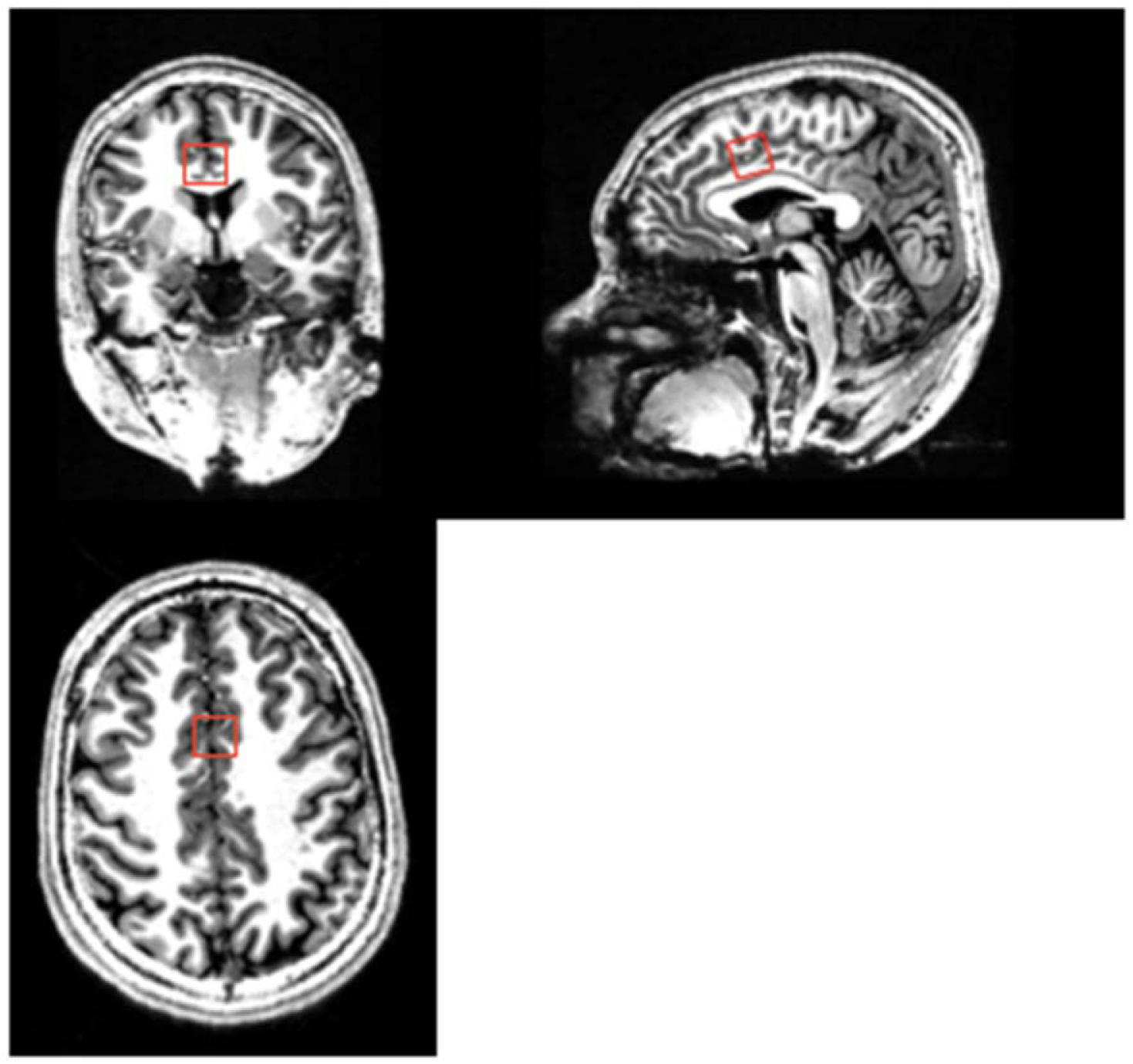
Dorsal Anterior Cingulate Cortex (ACC) voxel for MRS glutamate and glutathione estimation

### Statistical Analyses

All statistical tests were performed using IBM SPSS Statistics version 24. Differences in demographic and baseline factors between patients and controls were calculated using t-tests for continuous variables, and chi-square analyses for dichotomous variables. A linear regression analysis was used to assess the association between metabolites (glutamate and GSH), and both time to response, and social functioning (Hypotheses 1 and 3). Using ANOVA, we then compared glutamate and GSH measures among patients achieving remission at one month, no remission at one month, and healthy controls. Finally, Pearson correlation coefficients were used to assess the association between glutamate and GSH in patients and healthy controls. Differences in the magnitude of these correlations were then evaluated using Fisher’s r-to-Z transformation (Hypothesis 2).

## Results

### Patient Characteristics

37 patients completed baseline scanning. Of these, 27 met criteria for a schizophrenia spectrum disorder (SSD: schizophrenia, schizoaffective disorder, or schizophreniform disorder). Follow-up outcome data were not available for one female patient who was transferred to a different hospital shortly after scanning. In one male patient, time to response was not available due to irregular follow-up however, remission status at one month was obtained. Therefore, the final sample consisted of 26 patients with SSD, with time to response measures available for 25 patients (Table 1). See Supplement (SF1) for the representativeness of the sample. Based on Egerton et al^20^ (UK sample) reporting an effect size d=2.6 for ACC glutamate difference between 1-month remitters and non-remitters, we required a sample of at least 22 patients to demonstrate 50% of the reported effect (d=1.3), with 5% type 1 and 20% type 2 error rates.

**Table 1.**
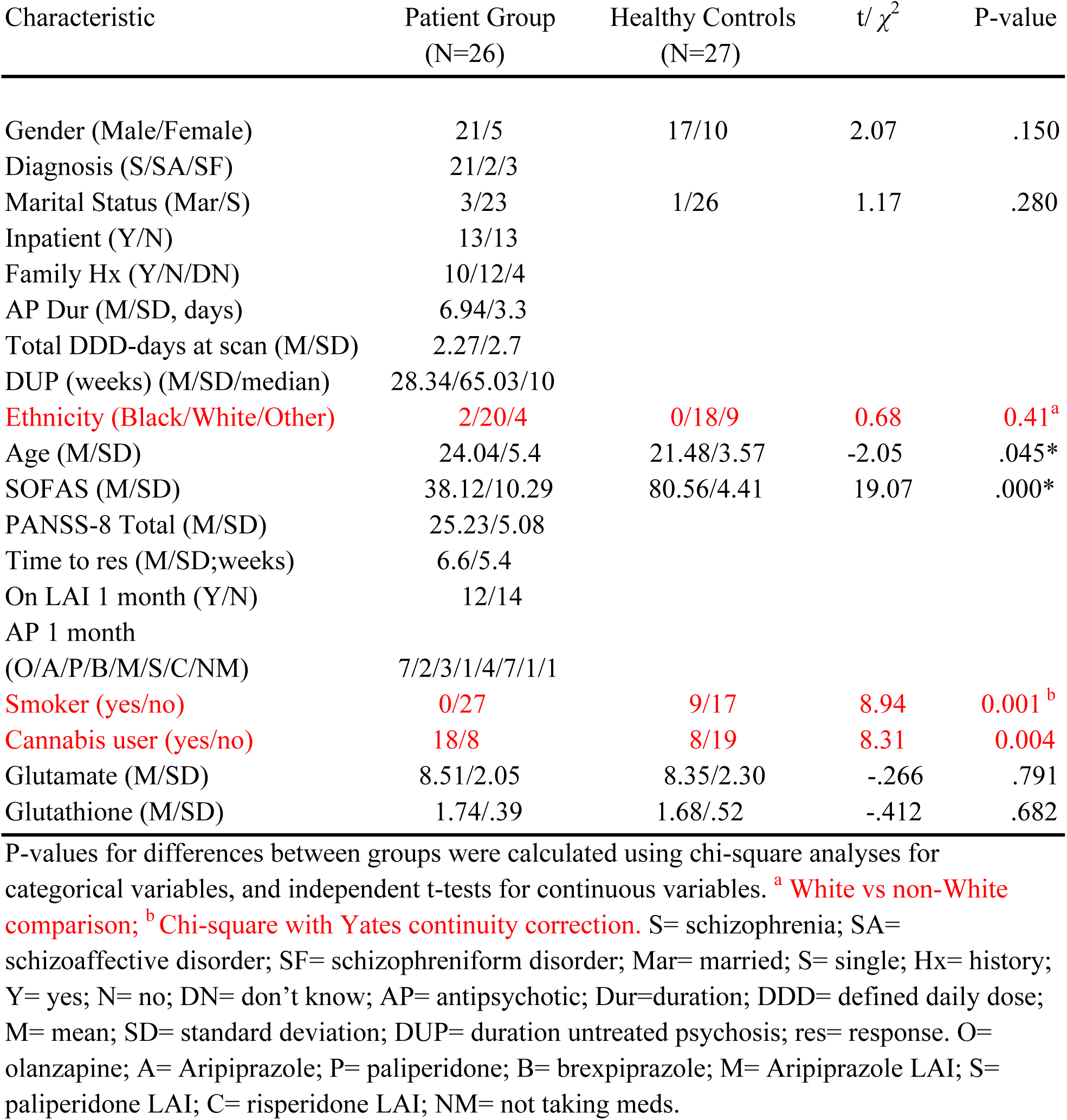
Sample Demographic and Clinical Characteristics.

| Characteristic | Patient Group<br>(N=26) | Healthy Controls<br>(N=27) | t/ $\chi^2$ | P-value |
| --- | --- | --- | --- | --- |
| Gender (Male/Female) | 21/5 | 17/10 | 2.07 | .150 |
| Diagnosis (S/SA/SF) | 21/2/3 |  |  |  |
| Marital Status (Mar/S) | 3/23 | 1/26 | 1.17 | .280 |
| Inpatient (Y/N) | 13/13 |  |  |  |
| Family Hx (Y/N/DN) | 10/12/4 |  |  |  |
| AP Dur (M/SD, days) | 6.94/3.3 |  |  |  |
| Total DDD-days at scan (M/SD) | 2.27/2.7 |  |  |  |
| DUP (weeks) (M/SD/median) | 28.34/65.03/10 |  |  |  |
| Ethnicity (Black/White/Other) | 2/20/4 | 0/18/9 | 0.68 | 0.41<sup>a</sup> |
| Age (M/SD) | 24.04/5.4 | 21.48/3.57 | -2.05 | .045* |
| SOFAS (M/SD) | 38.12/10.29 | 80.56/4.41 | 19.07 | .000* |
| PANSS-8 Total (M/SD) | 25.23/5.08 |  |  |  |
| Time to res (M/SD;weeks) | 6.6/5.4 |  |  |  |
| On LAI 1 month (Y/N) | 12/14 |  |  |  |
| AP 1 month<br>(O/A/P/B/M/S/C/NM) | 7/2/3/1/4/7/1/1 |  |  |  |
| Smoker (yes/no) | 0/27 | 9/17 | 8.94 | 0.001<sup>b</sup> |
| Cannabis user (yes/no) | 18/8 | 8/19 | 8.31 | 0.004 |
| Glutamate (M/SD) | 8.51/2.05 | 8.35/2.30 | -.266 | .791 |
| Glutathione (M/SD) | 1.74/.39 | 1.68/.52 | -.412 | .682 |

9 patients (34.6%) were antipsychotic naïve at the time of scanning, 5 patients were taking other psychotropic medications at the time of scanning as follows: 2 clonazepam, 1 lorazepam, 1 escitalopram and 1 sertraline. Of those who had already started antipsychotic treatment, (17; 65.4%), the median days of treatment was 6 (range of 3-12 days). The mean total defined daily dose-days (DDD X days on medication) for antipsychotic use was 2.27 days. At one month, 12 patients (46.15%) were taking a long acting injectable medication. In terms of cross-sectional remission, we observed the rates of 42.31% (n=11 of 26) a 1-month, 50% (n=13 of 26) at 3-months and 60% (n= 15 of 25) at 6 months. We did not stipulate cessation before scanning to avoid possible withdrawal effects and participants may have used nicotine on the day of scanning.

### ^1^H-MRS Data Quality

The mean glutamate CRLB percentages did not differ between healthy controls and patients (mean(SD) in % = 3.36 (1.02) in controls; 3.72(1.19) in FEP; t=1.16, p=0.25). Mean GSH CRLBs were (mean(SD) in % = 10.46(3.88) in controls; 11.47(4.92) in FEP; t=0.81, p=0.42). The percent coefficient of variation (%CV), calculated as the standard deviation divided by the mean of a sample, was 20.4% and 24.1% for healthy control and FEP glutamate measurements, respectively and 24.8% and 22.6% for healthy control and FEP GSH measurements, respectively (control vs FEP - p>0.6 for both metabolites). The average line width of the water-unsuppressed spectra did not differ between the 2 groups (mean(SD) = 7.62(1.17) in controls; 7.48(1.42) in FEP; t=0.39,p=0.7). The NAA peak-area signal-to-noise ratio was also not different (mean(SD) = 109.88 (18.37) in controls; 102.19 (24.53) in FEP; t=1.29,p=0.20), where the NAA peak-area SNR is defined as the ratio of the time-domain amplitude of the NAA CH_3_ singlet divided by the standard deviation of the noise measured in the last 32 points of the time-domain signal.

### GSH, Glutamate and Time to Response

Multiple regression analysis was used to test if GSH and glutamate significantly predicted the time taken by patients with FEP to respond to antipsychotic treatment. The results of the regression indicated the two predictors explained 31% of the variance (R2 =.0.31, F(2,24)=4.86, p=0.018). Higher levels of GSH predicted a shorter time to response (β = -0.65, p=0.017) while glutamate was not a significant predictor (β = 0.15, p=0.563) (see figure 2A). A very low level of multicollinearity was present (VIF = 1.98 for both GSH and glutamate). Results remained unchanged after controlling for age, sex, and daily dose of antipsychotics.

**Figure 2.**
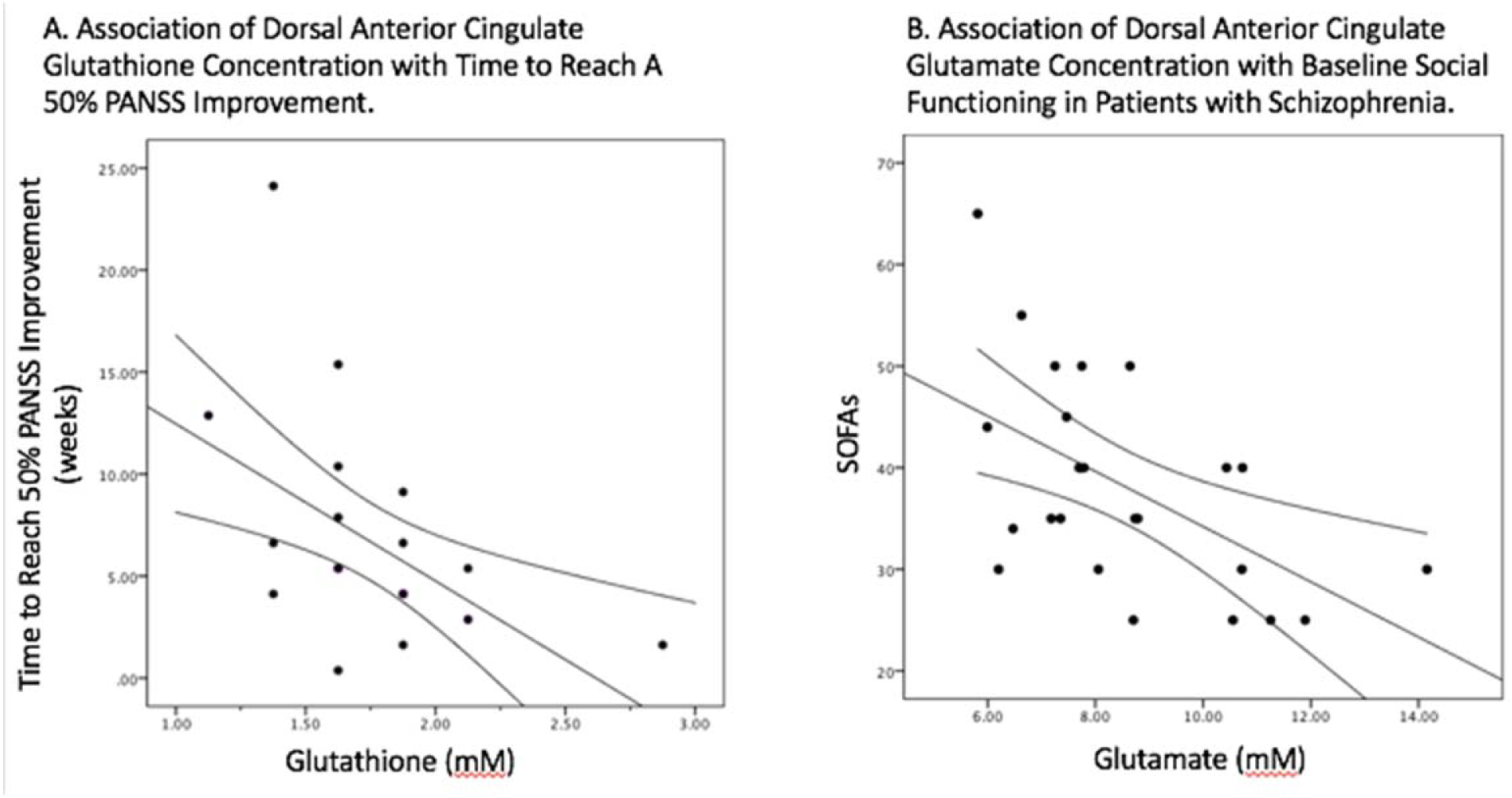
Association of Dorsal Anterior Cingulate Metabolites with Outcome Measures

### GSH, Glutamate and Social Functioning

Multiple regression analysis was used to test if GSH and glutamate significantly predicted the SOFAS scores in patients with FEP. The results of the regression indicated the two predictors explained 33% of the variance (R^2^ =.0.33, F(2,24)=5.33, p=0.013). Higher levels of glutamate predicted lower SOFAS scores (β = -0.70, p=0.008) while GSH was not a significant predictor (β = 0.22, p=0.376) (see figure 2B). A very low level of multicollinearity was present (VIF = 1.89 for both GSH and glutamate). Results remained unchanged after controlling for age, sex, and daily dose of antipsychotics.

### Correlations Between Metabolite Levels

The association between glutamate and GSH was tested using Pearson correlation coefficients. There was a positive association between levels of ACC glutamate and GSH in both healthy control subjects (*r*= .91, *p*< .001), and in patients with FEP (r= .69, *p*< .001). We then used Fisher’s r-to-z transformation to test the significance of difference between the correlations, and found that the correlation between glutamate and GSH was significantly weaker in patients compared to the healthy control subjects (Z= 2.26, *p*= .023) (see figure 3).

**Figure 3.**
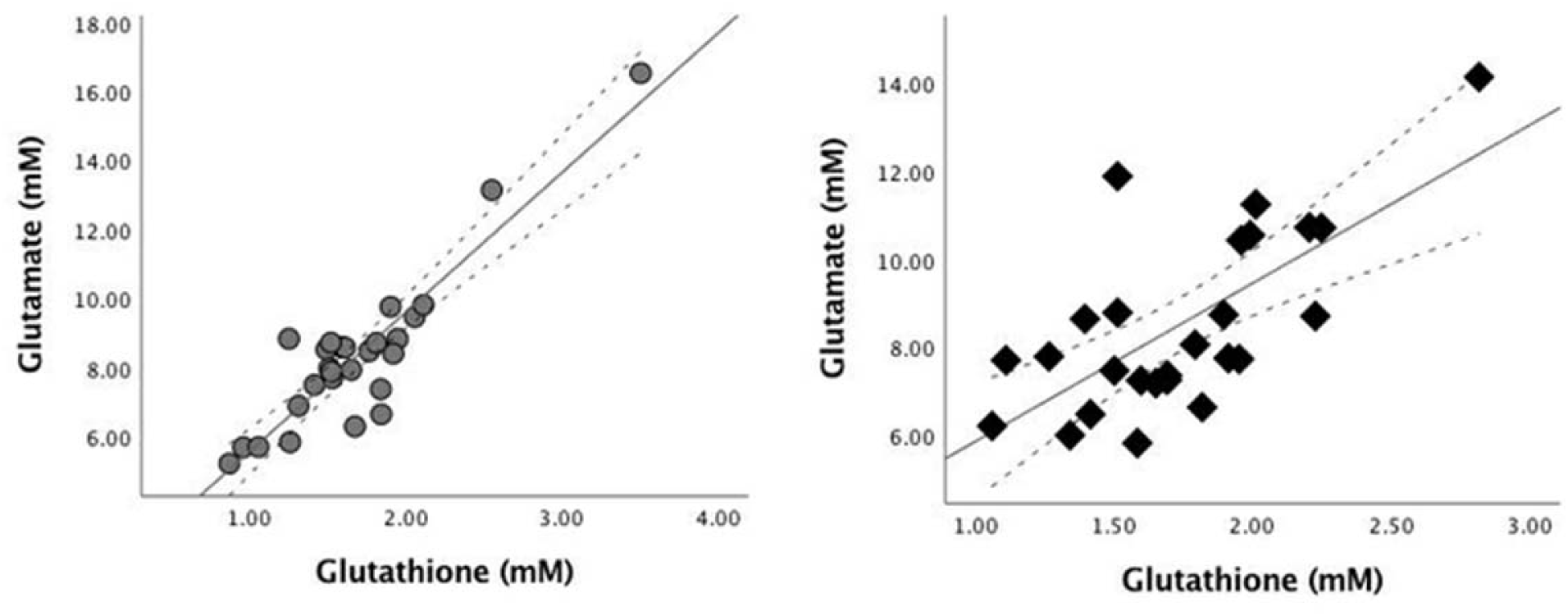
Correlation between Glutamate and Glutathione in Healthy Controls (left, circles) and Patients (right, diamonds). The metabolite concentrations are estimated in mM units.

### Group differences in GSH and Glutamate

One-way ANOVAs were conducted to evaluate the differences in metabolite levels among patients in remission or non-remission at one month and healthy control subjects. There were no significant difference between groups for glutamate (F(2,50)=.134, *p*=.875) or GSH (F(2,50)=.712, *p*=.496) (see Table 2). There were no significant differences between patients (as a single group) and controls on measures of glutamate (t(51)= -.266, *p*= .791) or GSH (t(51)= - .412, *p*=.682.

**Table 2.** ACC Glutamate and GSH levels in Patients in Remission, Not in Remission, and Healthy Controls

| Metabolite | All Patients (M/SD)<br>(N=26) | Remission (M/SD)<br>(N=11) | No Remission(M/SD)<br>(N=15) | HC (M/SD)<br>(N=27) |
| --- | --- | --- | --- | --- |
| Glutamate (mM) | 8.51/2.05 | 8.73/2.30 | 8.34/1.91 | 8.35/2.30 |
| GSH (mM) | 1.74/.39 | 1.85/.48 | 1.65/.30 | 1.68/.52 |

| Metabolite | All Patients (M/SD)<br>(N=26) | Remission (M/SD)<br>(N=11) | No Remission(M/SD)<br>(N=15) | HC (M/SD)<br>(N=27) |
| --- | --- | --- | --- | --- |
| Glutamate (mM) | 8.51/2.05 | 8.73/2.30 | 8.34/1.91 | 8.35/2.30 |
| GSH (mM) | 1.74/.39 | 1.85/.48 | 1.65/.30 | 1.68/.52 |

The effects of recreational substance use and types of antipsychotics are presented in the Supplement.

## Discussion

This is the first study to use ultra high-field 7T MRS to investigate the role of glutamate and GSH in early treatment response, and the first 7T MRS study on minimally medicated FEP subjects. A previous 7T MRS study included FEP subjects with an average of 55 weeks of antipsychotic exposure^54^, compared to 6 days of median exposure in our sample. A more recent study^23^ included FEP subjects with up to 2 years of illness duration, while we recruited all subjects during the acute first episode (mean SOFAS score of 38.1). We report 3 major findings (1) Patients with FEP with higher GSH levels in ACC show a rapid symptom reduction upon starting antipsychotic treatment (2) When compared to healthy controls, GSH levels in patients are dissociated from glutamate levels (3) Glutamate excess predicts the degree of Social and Occupational dysfunction seen at the time of presentation with FEP. Taken together, these results indicate that markers of cortical redox integrity influence the putative glutamatergic toxicity and early treatment response in psychosis.

Neither glutamate nor GSH were associated with binary remission status at one month. The lack of association is in contrast with the overall results reported by another study^17^, but consistent with the observation reported by 2 out of the 3 sites in that study. These differences can be attributed to methodological variations (the use of 7T spectra, more dorsal voxel placement in our study) as well as notable differences in the clinical samples (the use of injectables and the inclusion of both inpatients and outpatients, and the exclusion of patients with low adherence in our study). Egerton et al (2018)^17^ noted that higher glutamate levels correlated with greater symptom severity as well as poor functioning at baseline. Individuals, who are more severely ill, may be less likely to adhere to antipsychotic medications, with resulting ongoing symptom burden, and subsequent lack of remission. More recently, Iwata et al. (2018)^18^ found no differences in glutamate levels in the dorsal ACC between treatment resistant vs. responsive patients. Despite the above clinical and methodological differences, we observed a significant relationship between higher glutamate levels and lower social/occupational functioning, in line with Egerton et al, (2018)^17^ as well as prior observations from our centre^19, 20^. A low level of social functioning at FEP is reported to be a robust and independent predictor of later treatment^55^. This finding adds strength to the prevailing notion that glutamatergic excess plays a critical role in shaping the poor outcome trajectory in psychosis.

We found no significant differences in GSH levels between patients and healthy controls. This is not surprising, given that meta-analytic pooling of ACC GSH studies in schizophrenia reveal a small overall effect size^24^, suggesting the possibility of heterogeneity in the GSH levels and thus redox status among patients. Our results suggest that such heterogeneity may map onto antipsychotic responsiveness, resulting in the conflicting findings of GSH levels reported to date in schizophrenia^23^.

We found evidence that despite their significant within-group correlation, when compared to healthy controls, glutamate and GSH levels were less tightly correlated among patients with FEP. A similar dissociation was also reported by Xin et al, (2016) ^56^, especially among patients with a GCLC-risk genotype affecting GSH synthesis. These results indicate that in a subset of patients with FEP, concomitant GSH response fails to occur when demands arise due to glutamatergic excess. Such patients are likely to be vulnerable to neurotoxic damage^57^, poor treatment response, and greater functional decline as a result of unchecked neuronal/glial damage^58^. Interestingly in healthy controls, when glutamatergic synapses are active due a task demand, GSH levels appear to increase concomitantly with glutamate^59^.

There are several strengths to the current study. First, the use of a 7T MR scanner, with higher specificity in identifying the glutamate resonance^60^, is a considerable strength. Second, our increased use of LAI’s may have improved adherence rates in our sample. Third, patients were followed frequently (weekly) over the course of their early illness trajectory. Finally, our sample is unique in that we recruited patients before antipsychotic treatment was established. Limitations

Participants in our study were treated with various antipsychotics; we cannot rule out variations in response patterns based on differential medication treatment. Secondly, we could not recruit a completely antipsychotic-naïve sample for obvious ethical reasons. While it is possible that metabolite levels were affected by antipsychotic medication, our sample is comprised of the least-treated subjects of all MRS glutamate and GSH studies in schizophrenia reported to date (median treatment duration = 2.27 DDD-days). Animal studies have shown that neuroleptic administration in rodents, even over 2 days, can affect D2-receptor occupancy^61^; such rapid effects in human striatum may indirectly affect prefrontal glutamate levels, given the relationship between striatal dopamine and prefrontal glutamate^62^. While the effect of antipsychotics on cerebral GSH is still unknown, Ivanova et al, (2015)^63^ suggest that serum GSH is affected by typical but not atypical antipsychotics. None of our patients were exposed to typical antipsychotics at the time of scanning. Similarly, other psychotropic medications (although taken in small numbers, including 3 on benzodiazepines) may have influenced spectroscopic results. Henry et al, (2010)^64^ found no acute effect on glutamate in healthy volunteers treated acutely with benzodiazepines, though glutamine levels increased. See the Supplement for the statistical effect of adjustment for dose and type of antipsychotic medication.

A further limitation is that our spectroscopic analysis was limited to the dorsal ACC and did not include more anterior/ventral portions of the medial prefrontal cortex. We cannot completely rule out the effect of recreational substances on the observed results (see Supplement). One study^65^ found that ACC glutamate levels were decreased in individuals who used cannabis regularly, while these results were not replicated in another study^66^. To our knowledge, no studies have investigated the association of GSH with cannabis use and none have examined the effects of cannabis on metabolite levels specifically in a FEP sample. Finally, our patient sample consisted primarily of males, limiting generalization of the results.

A promising implication is that interventions that increase GSH levels early in FEP may have the potential to alter the prognostic trajectory of psychosis (See Supplement -Translational Relevance for further details). A prospective sequential treatment trial^67^ in first episode patients has indicated that merely switching antipsychotics may not boost treatment response in early non-responders, and second level treatments such as clozapine may be warranted even before the conventional clinical threshold of Treatment Resistant Schizophrenia (i.e. 2 treatment failures) is met. While early non-response is considered to be an indicator of later non-response and subsequent treatment resistance in schizophrenia^68^, to our knowledge, the association between early non-response in first-episode samples and later sequential treatment failures and the status of conventionally defined TRS is yet to be established. One of the challenges in this regard is the high degree of responsiveness to treatment seen in first-episode patients^69^ (also observed in the current study), compared to those with acute exacerbation of chronic schizophrenia^68^. In this context, caution is warranted in extrapolating the physiological correlates of early treatment response as indicators of the emergence of categorical treatment resistance at later stages of schizophrenia. Given that GSH levels have a significant impact on the speed of response, we urge further experimental trials that manipulate GSH levels to observe the predicted gain in trajectory of treatment outcomes in FEP.

Preliminary results have demonstrated that NAC, a GSH precursor, may be beneficial in psychotic disorders^70^. NAC has been shown to be efficacious in reducing the symptom burden^71^, especially negative^72^ and cognitive symptoms^73^, and has the potential to alleviate treatment resistance in schizophrenia^74^. Our results suggest that treatments such as NAC may be efficacious particularly in patients who demonstrate an early poor response to antipsychotic medication, as they are likely to have a lower ability to synthesize GSH in response to glutamatergic excess. While MRS indices are indirect measures of tissue metabolite concentrations^75^, given the evidence that oral NAC administration in patients with schizophrenia increases GSH content in the ACC^70^, we consider MRS as a viable tool for translational investigations into the redox abnormalities of schizophrenia. More speculatively, we suggest that the association of ACC GSH levels at baseline and eventual clozapine-eligibility would be worth investigating in the future, given the lack of objective predictors of clozapine requirement in schizophrenia.

## Acknowledgements

We thank Dr. Joe Gati, Mr. Trevor Szekeres, Dr. Tushar Das and Dr. Ali Khan for their assistance in data acquisition and archiving. We thank Dr. William Pavlovsky for consultations on clinical radiological queries. We thank Drs. Raj Harricharan, Julie Richard, Priya Subramaniam and Hooman Ganjavi and all staff members of the PEPP London team for their assistance in patient recruitment and supporting clinical care. We gratefully acknowledge the participants and their family members for their contributions. The MRS pulse sequence package used was developed from a source code provided by Uzay E. Emir, PhD, and Dinesh K. Deelchand, PhD, Center for Magnetic Resonance Research (CMRR), Minneapolis, MN. It is based on original work by Gülin Öz and Ivan Tkáč [1. Öz G, Tkáč I (2011) Short-echo, single-shot, full-intensity 1H MRS for neurochemical profiling at 4T: Validation in the cerebellum and brainstem, Magn Reson Med, 65(4):901-10]. Our special thanks to Uzay E. Emir, PhD, Dinesh K. Deelchand, PhD, Gülin Öz, PhD, Ivan Tkáč, PhD, and Edward Auerbach, PhD, for their help with sequence testing and for valuable technical discussions and to Dennis W. J. Klomp, PhD, and Vincent O. Boer, PhD, for providing the asymmetric excitation pulse used in the 7T version of the sequence. We also acknowledge CFREF BrainsCAN funding that we received providing a reduced hourly rate for the MRI scans.

## References

1. Schennach-Wolff R, Seemuller FH, Mayr A, Maier W, Klingberg S, Heuser I, et al. An early improvement threshold to predict response and remission in first-episode schizophrenia. Br J Psychiatry 2010; 196:460–466.

2. Stauffer VL, Case M, Kinon BJ, Conley R, Ascher-Svanum H, Kollack-Walker S, et al. Early response to antipsychotic therapy as a clinical marker of subsequent response in the treatment of patients with first-episode psychosis. Psychiatry Res 2011; 187(1-2):42–48.

3. Emsley R, Osthulzen PP, Kidd M, Koen L, Niehaus DJ, Turner HJ. Remission in first-episode psychosis: predictor variables and symptom improvement patterns. J Clin Psychiatry 2006; 67(11):1707–1712.

4. Kinon BJ, Chen L, Ascher-Svanum H, Stauffer VL, Kollack-Walker S, Sniadecki JL, et al. Predicting response to atypical antipsychotics based on early response in the treatment of schizophrenia. Schizophr Res 2008; 102(1-3):230–240.

5. Lindenmayer JP. Treatment refractory schizophrenia. Psychiatr Q 2000; 71:373–384.

6. Lally J, Ajnakina O, Di Forti M, Trotta A, Demjaha A, Kolliakou A, et al. Two distinct patterns of treatment resistance in first-episode schizophrenia-spectrum psychoses. Psychol Med 2016; 46:3231–3240.

7. Demjaha A, Lappin JM, Stahl D, Patel MX, MacCabe JH, Howes OD et al. (2017): Antipsychotic treatment resistance in first-episode psychosis: prevalence, subtypes and predictors. Psychol Med 2017; 47:1981–1989.

8. Poels EMP, Kegeles LS, Kantrowitz JT, Javitt DC, Lieberman JA, Abi-Dargham A, et al. Glutamatergic abnormalities in schizophrenia: A review of proton MRS findings. Schizophr Res 2014; 152:325–332

9. Merritt K, Egerton A, Kempton MJ, Taylor MJ, McGuire PK. Nature of glutamate alterations in schizophrenia: a meta-analysis of proton magnetic resonance spectroscopy studies. JAMA Psychiatry 2016; 73(7):665–674.

10. Jauhar S, McCutcheon R, Borgan F, Veronese M, Nour M, Pepper F, et al. The relationship between cortical glutamate and striatal dopamine in first-episode psychosis: A cross-sectional multimodal PET and magnetic resonance spectroscopy imaging study. The Lancet Psychiatry 2018;5:816–823.

11. Abi-Dargham A, Rodenhiser J, Printz D, Zea-Ponce Y, Gil R, Kegeles L, et al. Increased baseline occupancy of D2 receptors by dopamine in schizophrenia. Proceedings of the National Academy of Sciences, USA 2000;97:8104–8109.

12. Howes O, McCutcheon R, Stone J. Glutamate and dopamine in schizophrenia: an update for the 21st century. J Psychopharmacol Oxf Engl 2015;29:97–115. https://doi.org/10.1177/0269881114563634

13. Mouchlianitis E, Bloomfield MAP, Law V, Beck K, Selvaraj S, Naresh Rasquinha, et al. Treatment-resistant schizophrenia patients show elevated anterior cingulate cortex glutamate compared to treatment-responsive. Schizophr Bull 2016; 42:744–752

14. Szulc A, Konarzewska B, Galinska-Skok B, Lazarczyk J, Waszkiewicz N, Tarasow E, et al. Proton magnetic resonance spectroscopy measures related to short-term symptomatic outcome in chronic schizophrenia. Neurosci Lett 2013; 547:37–41.

15. Demjaha A, Egerton A, Murray RM, et al. Antipsychotic treatment resistance in schizophrenia associated with elevated glutamate levels but normal dopamine function. Biol Psychiatry 2014; 75:11–13.

16. Egerton A, Brugger S, Raffin M, Barker GJ, Lythgoe GJ, McGuire PK et al. Anterior cingulate glutamate levels related to clinical status following treatment in first-episode schizophrenia. Neuropsychopharmacology 2012; 37:2515–2521.

17. Egerton A, Broberg BV, Van Haren N, Merrit K, Barker GJ, Lythcoe DJ, et al. Response to initial antipsychotic treatment in first episode psychosis is related to anterior cingulate glutamate levels: A multicenter 1H-MRS study (OPTiMiSE). Mol Psychiatry 2018 (in press).

18. Iwata Y, Nakajima S, Plitman E, Mihashi Y, Caravaggio F, Chung JK, et al. Neurometabolite levels in antipsychotic-naïve/free patients with schizophrenia: A systematic review and meta-analysis of ^1^H-MRS studies. Progress in Neuro-Psychopharmacology and Biological Psychiatry 2018; 86:340–352.

19. Aoyoma N, Theberge J, Drost DJ, Manchanda R, Northcott S, Neufeld RW, et al. Grey matter and social functioning correlates of glutamatergic metabolite loss in schizophrenia. Br J Psychiatry 2011; 198(6):448–456.

20. Dempster K, Norman R, Theberge J, Densmore M, Schaefer B, Williamson P. Glutamatergic metabolite correlations with neuropsychological tests in first episode schizophrenia. Psychiatry Res Neuroimaging 2015; 233:180–185.

21. Mahadik, S.P. & Mukherjee, S. Free radical pathology and antioxidant defense in schizophrenia ± a review. Schizophr Res 1996; 19:1–17.

22. Do KQ, Cabungcal JH, Frank A, Steullet P, Cuenod M. Redox dysregulation, neurodevelopment, and schizophrenia. Curr Opin Neuobiol 2009; 19(2):220–230.

23. Wang AM, Pradhan S, Coughlin JM, Trivedi A, DubBois SL, Crawford JL, et al. Assessing brain metabolism with 7-T proton magnetic resonance spectroscopy in patients with first-episode psychosis. JAMA Psychiatry 2019; (in press).

24. Das TK, Javadzadeh A, Dey A, Sebasan P, Theberge J, Radua J, Palaniyappan P. Antioxidant defense in schizophrenia and bipolar disorder: A meta-analysis of MRS studies of anterior cingulate glutathione. Prog Neuropsychopharmacol Biol Psychiatry 2018; (in press).

25. Kumar J, Liddle EB, Fernandes CC, Palaniyappan L, Hall EL, Robson SE, et al. Glutathione and Glutamate in schizophrenia: A 7T MRS Study. Mol Psychiatry 2018 (in press).

26. Klauser P, Xin L, Fournier M, Griffa A, Cleusix M, Jenni R, et al. N-acetyl-cysteine add-on treatment leads to an improvement of fornix white matter integrity in early psychosis. Schizophr Bull 2018; 44(1):5133–5134.

27. Sedlak TW, Paul BD, Parker GM, Hester LD, Snowman AM, Taniguchi Y, et al. The glutathione cycle shapes synaptic glutamate activity. PNAS 2019; (in press).

28. Volterra A, Trotti D, Tromba C, Floridi S, Racagni G. Glutamate uptake inhibition by oxygen free radicals in rat cortical astrocytes. J Neurosci 1994; 14:2924–2932.

29. Frade J, Pope S, Schmidt M, Dringen R, Barbosa R, Pacock J, et al. Glutamate induces release of glutathione from cultured rat astrocytes-a possible neuroprotective mechanism. Journal of Neurochemistry 2008; 105(4):1144–1152.

30. Shih AY, Erb H, Sun X, Toda S, Kallivas PW, Murphy TH. Cystine/glutamate exchange modulates glutathione supply for neuroprotection from oxidative stress and cell proliferation. J Neurosci 2006; 26(41):10514–10523.

31. Fournier M, Monin A, Ferrari C, Baumann PS, Conus P, Do K. Implication of the glutamate-cystine antiporter xCT in schizophrenia cases linked to impaired GSH synthesis. NPJ Schizophr 2017; 3:31.

32. Plitman E, Patel R, Chung JK, Pipitone J, Chavez S, Reyes-Madrigal F, et al. Glutamatergic Metabolites, Volume, and Cortical Thickness in Antipsychotic-Naïve Patients with First Episode Psychosis: Implications for Excitotoxicity. Neuropsychopharmacology 2016; 41:2606–2613.

33. Leckman JF, Sholomskas D, Thompson WD, Belanger A, Weissman MM. Best estimate of lifetime psychiatric diagnosis: A methodological study. Arch Gen Psychiatry 1982; 39:879–883.

34. First MB, Williams JBW, Karg RS, Spitzer RL. Structured Clinical Interview for DSM-5 Disorders, Clinician Version (SCID-5-CV). Arlington, VA, American Psychiatric Association, 2016.

35. Remington G, Addington D, Honer W, Ismail Z, Raedler T, Teehan M. Guidelines for the Pharmacotherapy of Schizophrenia in Adults. Can J Psychiatry 2017; 62:604–616.

36. American Psychiatric Association. (2013). Diagnostic and statistical manual of mental disorders (5^th^ ed.). Arlington, VA:American Psychiatric Publishing

37. Norman R, Anderson K, MacDougall A, Manchanda R, Harricharan R, Subramanian P, Richard J, Northcott S. Stability of outcomes after 5 years of treatment in an early intervention program. Early intervention in Psychiatry, 2018; 12:720–725.

38. Harvey PD, Bellack AS. Toward a terminology of functional recovery in schizophrenia: Is functional remission a viable concept? Schizophr Bull 2009; 35:300–306.

39. Andreasen NC, Carpenter Jr WT, Kane JM, Lasser RA, Marder SR, Weinberger DR. Remission in schizophrenia: proposed criteria and rationale for consensus. Am J Psychiatry 2005; 162:441–449.

40. Lin CH, Lin HS, Kuo CC, Wang FC, Huang YH. Early Improvement in PANSS-30, PANSS-8 and PANSS-6 scores predicts ultimate response and remission during acute treatment of schizophrenia. Acta Psychiatrica Scandinavica 2018; 137:98–108.

41. Kay SR, Fiszbein A, Opler LA. 1987. The Positive and Negative Syndrome Scale (PANSS) for schizophrenia. Schizophr Bull 1987; 13:261–276.

42. Leucht, Stefan, Romain Beitinger, and Werner Kissling. On the concept of remission in schizophrenia. Psychopharmacology 2007; 194: 453–461.

43. Haro JM, Kamath SA, Ochoa S, Novick D, Rele K, Fargas A, et al. The Clinical Global Impression-Schizophrenia scale: a simple instrument to measure the diversity of symptoms present in schizophrenia. Acta Psychiatr Scand Suppl 2003; 416:16–23.

44. Correll C, Kishimoto T, Nielsen J, Kane J. Quantifying clinical relevance in the treatment of schizophrenia. Clin Ther 2011; 33:B16–B39.

45. Obermeier M, Mayr A, Schennach-Wolff R, Seamuller F, Moller H-J, Riedel M. Should the PANSS be rescaled? Schizophr Bull 2010; 36:455–460.

46. Goldman HH, Skodol AE, Lave TR. Revising axis V for DSM-IV: a review of measures of social functioning. Am J Psychiatry 1992; 149:1148–1156.

47. Cassidy CM, Rabinovitch M, Schmitz N, Joober R, Malla A. A comparison study of multiple measures of adherence to antipsychotic medication in first-episode psychosis. J Clin Pharmacol 2010; 30:64–67.

48. Tkáć I, Gruetter R. Methodology of 1 H NMR spectroscopy of the human brain at very high magnetic fields. Appl Magn Reson 2005; 29:139–157.

49. Near J, Edden R, Evans CJ, Paquin R, Harris A, Jezzard P. Frequency and phase drift correction of magnetic resonance spectroscopy data by spectral registration in the time domain. Magn Reson Med 2015; 73:44–50.

50. Bartha R, Drost DJ, Menon RS, Williamson PC. Spectroscopic lineshape correction by QUECC: Combined QUALITY deconvolution and Eddy current correction. Magn Reson Med 2000; 44:641–645.

51. Bartha R, Drost DJ, Williamson PC. Factors affecting the quantification of short echo in vivo 1H MR spectra: prior knowledge, peak elimination, and filtering. NMR Biomed 1999; 12:205–216.

52. Wong D, Schranz A, Bartha R. Optimized in vivo brain glutamate measurement using long-echo time semi-LASER at 7 T. NMR in Biomedicine 2018:e4002 https://doi.org/10.1002/nbm.4002

53. Stanley JA, Drost DJ, Williamson PC, Thompson RT. The use of a priori knowledge to quantify short echo in vivo 1H MR spectra. Magn Reson Med 1995; 34:17–24.

54. Reid MA, Salibi N, White DM, Gawne TJ, Denney TS, Lahti AC. 7T Proton Magnetic Spectroscopy of the Anterior Cingulate Cortex in First Episode Schizophrenia. Schizophr Bull 2018 (in press).

55. Horsdal HT, Wimberley T, Kohler-Forsberg O, Baandrup L, Gasse C. Association between global functioning at first schizophrenia diagnosis and treatment resistance. Early Intervention in Psychiatry 2018; 12:1198–1202.

56. Xin L, Mekle R, Fournier M, Baumann PS, Ferrari C, Alameda L, et al. Genetic polymorphism associated prefrontal glutathione and its coupling with brain glutamate and peripheral redox status in early psychosis. Schizophr Bull 2016; 42(5):1185–1196.

57. Hulshoff Pol HE, Kahn RS. What happens after the first episode? A review of progressive brain changes in chronically ill patients with schizophrenia. Schizophr Bull 2008; 34:354–366.

58. Kantrowitz JT, Javitt DC. N-methyl-d-aspartate (NMDA) receptor dysfunction or dysregulation: the final common pathway on the road to schizophrenia? Brain Res Bull 2010; 83:108–121.

59. Lin Y, Stephenson MC, Xin L, Napolitano A, Morris PG. Investigating the metabolic changes due to visual stimulation using functional proton magnetic resonance spectroscopy at 7T. J Cereb Blood Flow Metab 2012; 32:1484–1495.

60. Pradhan S, Bonekamp S, Gillen JS, Rowland LM, Wijtenburg SA, Edden RAE, et al. Comparison of single voxel brain MRS AT 3 T and 7 T using 32-channel head coils. Magn Reson Imaging 2015; 33:1013–1018.

61. Samaha AN, Seeman P, Stewart J, Rajabi H, Kapur S. “Breakthrough” dopamine supersensitivity during ongoing antipsychotic treatment leads to treatment failure overtime. J Neurosci 2007; 27:2979–2986.

62. Jauhar S, McCutcheon R, Borgan F, Veronese M, Nour M, Pepper F, et al. The relationship between cortical glutamate and striatal dopamine in first-episode psychosis: A cross-sectional multimodal PET and magnetic resonance spectroscopy imaging study. The Lancet Psychiatry 2018;5:816–823.

63. Ivanova SA, Smirnova LP, Shchigoreva YG, Semke AV, Bokhan NV. Serum glutathione in patients with schizophrenia in dynamics of antipsychotic therapy. Bull Exp Biol Med 2015; 160(2):283–285.

64. Henry ME, Jensen JE, Licata SC, Ravichandran C, Butman ML, Shanahan M, et al. The acute and late CNS glutamine response to benzodiazepine challenge: a pilot pharmacokinetic study using proton magnetic resonance spectroscopy. Psychiatry Res 2010;184:171–176.

65. Prescot AP, Renshaw PF, Yurgelun-Todd DA. γ-Amino butyric acid and glutamate abnormalities in adolescent chronic marijuana smokers. Drug Alcohol Depend 2013;129:232–239.

66. Sung Y-H, Carey PD, Stein DJ, Ferrett HL, Spottiswoode BS, Renshaw PF, et al. Decreased frontal N-acetylaspartate levels in adolescents concurrently using both methamphetamine and marijuana. Behav Brain Res 2013;246:154–161.

67. Kahn RS, Rossum Inge Winter van, Leucht S, McGuire P, Lewis SW, Leboyer M, et al. Amisulpride and olanzapine followed by open-label treatment with clozapine in first-episode schizophrenia and schizophreniform disorder (OPTiMiSE): a three-phase switching study. Lancet Psychiatry 2018;5:797–807.

68. Carbon M, Correll CU. Clinical predictors of therapeutic response to antipsychotics in schizophrenia. Dialogues Clin Neurosci 2014; 16:505–524.

69. Robinson DG, Woerner MG, Alvir J, Ma J, Geisler S, Koreen A, et al. Predictors of Treatment Response From a First Episode of Schizophrenia or Schizoaffective Disorder. Am J Psychiatry 1999;156:544–549.

70. Conus P, Seidman LJ, Fournier M, Xin L, Cleusix M, Baumann PS, et al. N-acetylcysteine in a double-blind randomized placebo-controlled trial: toward biomarker-guided treatment in early psychosis. Schizophr Bull 2018; 44:317–327.

71. Zheng W, Zhang Q-E, Cai D-B, Yang X-H, Qiu Y, Ungvari GS, et al. N-acetylcysteine for major mental disorders: a systematic review and meta-analysis of randomized controlled trials. Acta Psychiatr Scand 2018; 137:391–400.

72. Breier A, Leffick E, Hummer TA, Vohs JL, Yang Z, Mehdiyoun NF, et al. Effects of 12-month double-blind N-acetyl cysteine on symptoms, cognition and brain morphology in early phase schizophrenia-spectrum disorders. Schizophr Res 2018; 199:395–492.

73. Rapado-Castro M, Dod S, Bush AI, Skvarc DR, On ZX, Berk M, et al. Dean OM. Cognitive effects of adjunctive N-acetyl cysteine in psychosis. Psychol Med 2017; 47:866–876.

74. Bulut M, Savas HA, Altindag A, Virit O, Dalkilic A. Beneficial effects of N-acetylcysteine in treatment resistant schizophrenia. World J Biol Psychiatry 2009; 10:626–628.

75. Schwerk A, Alves FDS, Pouwels PJW, van Amelsvoort T. Metabolic alterations associated with schizophrenia: a critical evaluation of proton magnetic resonance spectroscopy studies. J Neurochem 2014;128:1–87.

