## Supplement for "Early Treatment Response in First Episode Psychosis: A 7-Tesla Magnetic Resonance Spectroscopic Study of Glutathione and Glutamate"

Representativeness of the sample	1

MRS data acquisition and quality	2

Clinical relevance	8

The use of PANSS-8 to assess clinical outcome	9

Antipsychotics, clinical outcome and MRS measures	10

Substance use, clinical outcome and MRS measures	11

References	13

### Representativeness of the sample

**Supplementary Figure 1. Flowchart of Patient Recruitment and Formation of Study Cohort.**

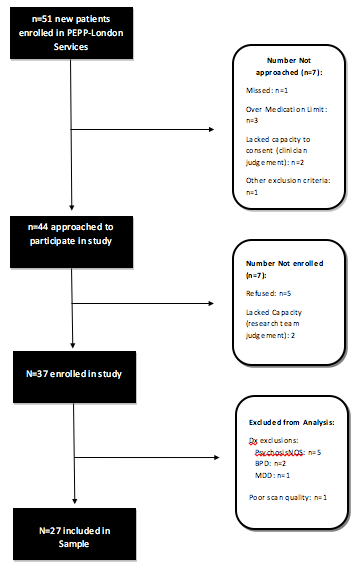

### MRS data acquisition and quality

**Supplementary Figure 2. Example of Spectral Fit.** The figure below is an example of a spectral fit using the fitMAN software. The yellow line represents the raw data, the red line represents the fitted data, and the teal line represents the residual.

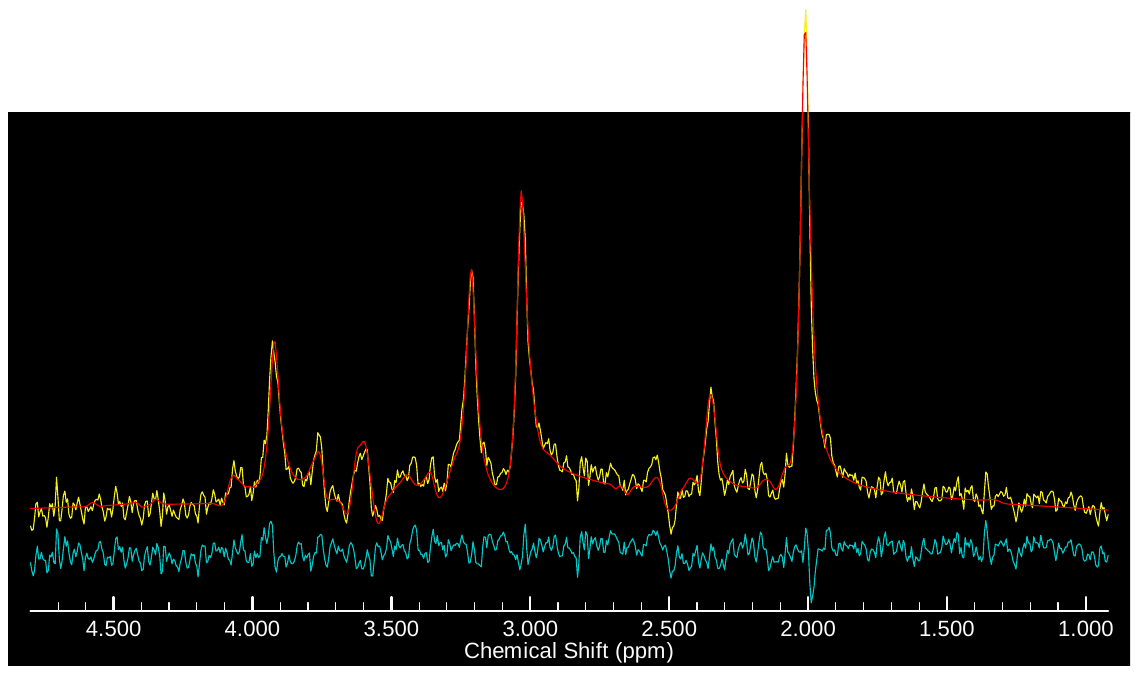

**Supplementary Figure 3. Metabolite breakdown from spectral fit:** The figure below depicts metabolite components of interest used in the Barstool software. The first line represents the residual between the raw and the fitted data. The lines marked as Data/Fit represent the raw (grey line) and the fitted data (superimposed black line). The grey line marked ‘Others’ represents the sum of 14 metabolites used in the fitting model; the next four grey lines represent the separated glutathione, glutamate, glutamine and GABA contributions toward the total fit data.

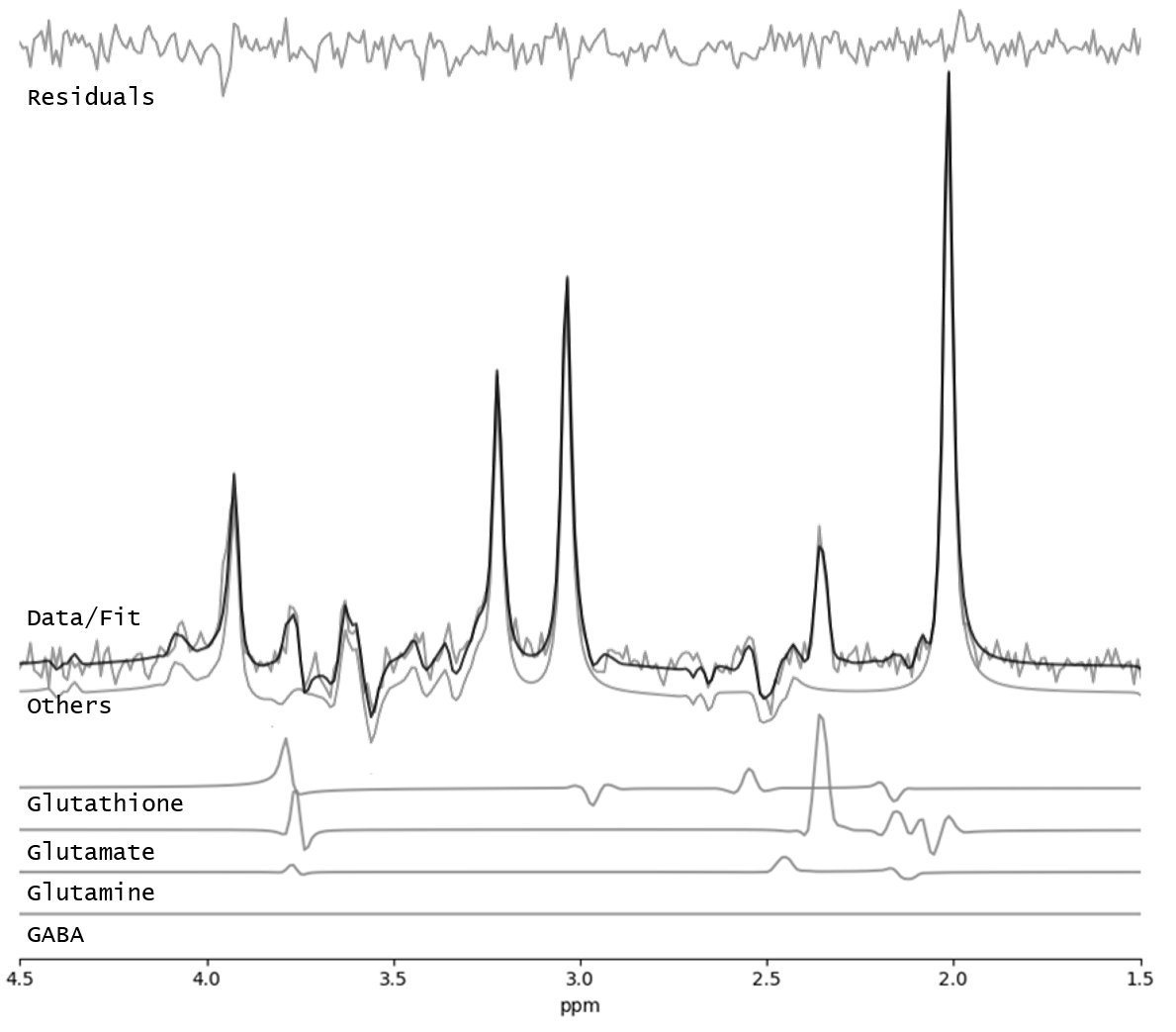

**Anatomical Voxel Location**

We used the following anatomical approach for placing the MRS voxel relative to known anatomical landmarks. The center was placed on the junction of the right cingulate sulcus with the paracentral sulcus (angled at the antero-posterior line, tangential to the corpus callosum), with the base of the voxel resting on the grey matter fold immediately above corpus callosum in the sagittal sections. In transverse sections, the voxel was centered on the interhemispheric fissure.

The dorsal anterior cingulate voxel was chosen as this region has been shown to be an integral part of the Salience Network^1,2^ and contributes to the performance of tasks that require selective attention^3^. Both Salience Network dysfunction 4,5 and selective attentional impairment^6,7^ have been demonstrated reliably in psychosis. Our dorsal ACC voxel placement also overlapped with several recent 7T MRS studies in psychosis^8,9^ (Supplementary Figure 4). Using meta-analytical synthesis of functional connectivity maps in Neurosynth (<http://neurosynth.org>), we demonstrate that the center of our voxel forms part of the Salience Network of the brain (see below for the Supplementary Figure 5).

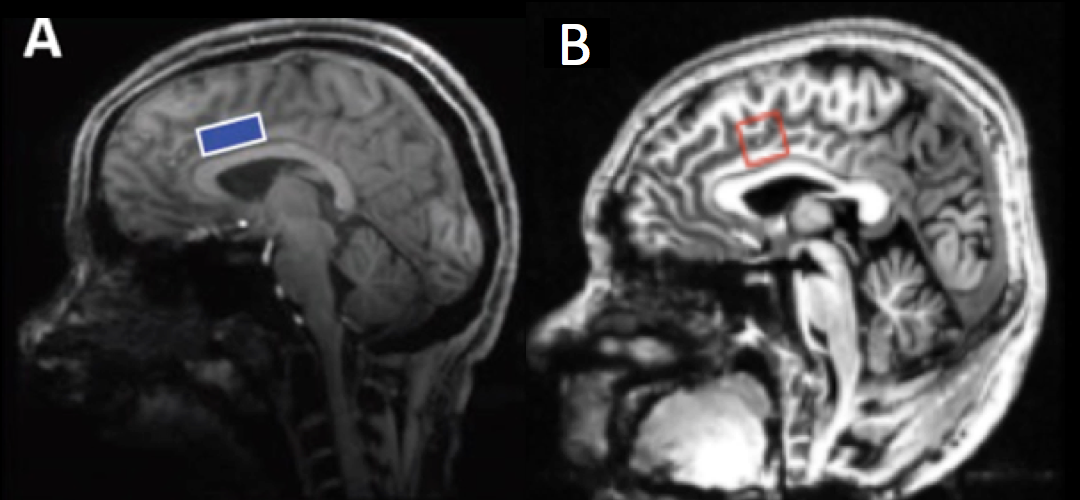

**Supplementary Figure 4.** Left (A): The MRS voxel used by Reid et al (2018) and Overbeek et al (2018) (Figure adapted from Overbeek et al (2018), supplementary material). Right (B): The MRS voxel used in the current study. Both images display a mid-sagittal slice from anatomical images acquired concurrently.

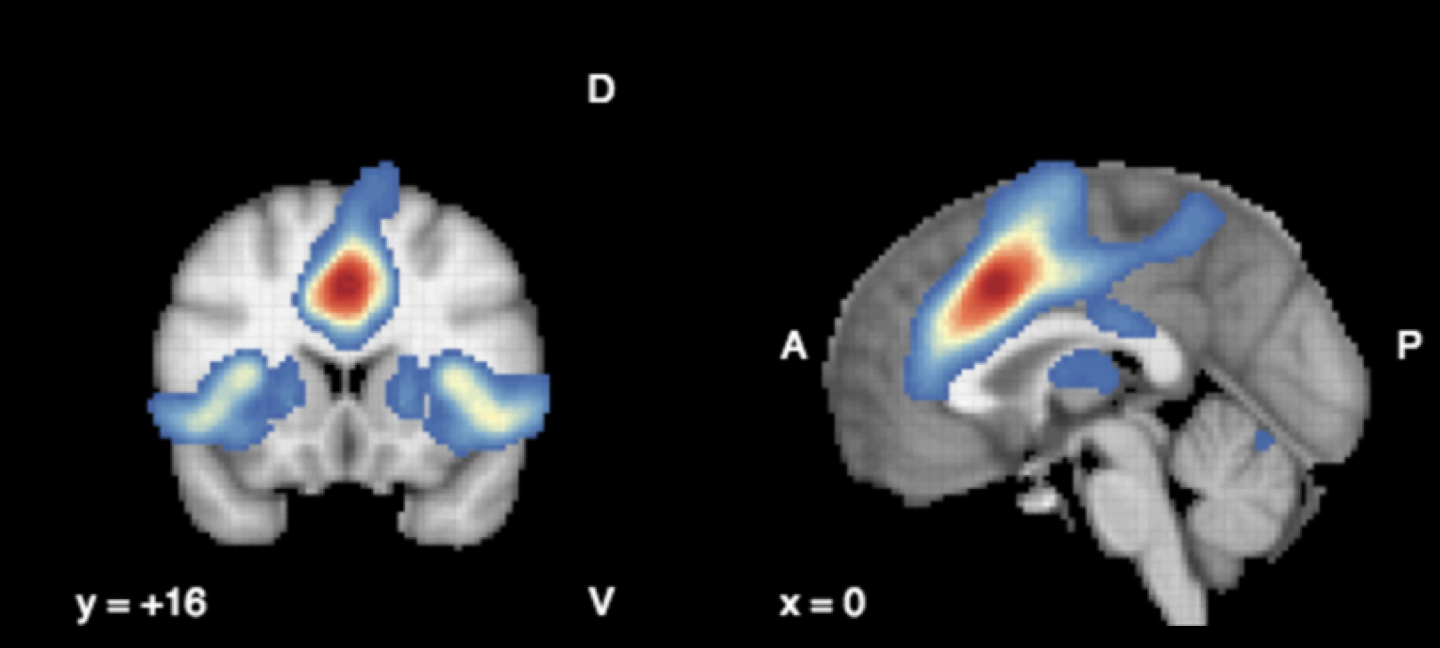

**Supplementary Figure 5.** The anatomical center of the dorsal ACC voxel used in the current study is functionally connected with the Salience Network of the brain (bilateral anterior insula and dorsal striatum). We used the anatomical center to query neurosynth database of functional connectivity (shown here) and functional coactivation (overlapping regions, not shown). D=Dorsal; V=Ventral; A=Anterior; P=Posterior.

**The use of CRLB as a quality measure**

Using relative Cramer-Rao Lower Bound (CRLB) as a quality measure to discard MRS data has been controversial. Kreis (2016)^10^ recommends that “relative CRLB should not be used to eliminate bad MRS data—certainly not as a sole criterion—because they may just reflect low levels of the measured quantity”. For metabolites with low concentration such as GSH, which are expected to be reduced in patients, this is of special concern, as this may systematically bias towards null results in group comparison. To study this further, we first established that the group level CRLBs were not different between patients and controls for glutamate and GSH (as reported in the manuscript). We then related CRLB values to the estimated metabolite concentrations, and found a strong negative correlation between the measured quantity of GSH and the reported CRLB across all of the subjects included in this study (r= -0.36, p=0.01). In line with this view, several recent studies have rejected the use of CRLB bounds or used absolute CRLB limits that indicate failure of model fit^11^ while others have used a liberal 30% cut-off for metabolites with smaller concentration^12^

In our data, we did not find any subject with CRLB>28% for GSH and CRLB>7.6% for glutamate. When we removed the 1 subject with GSH CRLB above 25%, none of our results were altered. Multiple regression analysis to test if GSH and glutamate significantly predicted the time taken by patients with FEP to respond to antipsychotic treatment continued to indicate that the two predictors GSH and glutamate explained 35% of the variance (F=5.62, p=0.011). Higher levels of GSH predicted a shorter time to response (β = -0.83, p=0.01) while glutamate was not a significant predictor (β = 0.35, p=0.25). Multiple regression analysis to test if GSH and glutamate significantly predicted the SOFAS continued to be significant (R^2^ =.0,28 F=3.99, p=0.034). Higher levels of glutamate predicted lower SOFAS scores (β = -0.68, p=0.03) while GSH was not a significant predictor (β = 0.22, p=0.47). We continued to see no significant differences between patients and controls on measures of glutamate (t= -.06, *p*= .95) or GSH (t(50)= -.22, *p*=.83).

 The echo time we used was selected purposely to improve glutamate quantification, while achieving valid and reliable GSH quantification. With respect to the validity of the GSH measure, it is worth noting that if we were fitting noise instead of real GSH peaks, the %CV would be numerically close to %CRLB.  In our sample, we observed a very different pattern (i.e., %CVs are in the ~20% for both GSH and Glu while the %CRLB are ~3% for Glu and 10% for GSH).   Previous work in our centre^13^ has demonstrated that while GSH was correlated with glutamine at 1.5T (likely due to overlap), this was not the case at 7T (much less overlap). Taken together, these observations support the validity of the GSH and Glu signal measured in the current study.

**The effect of outliers on the regression analyses**

We observed an outlier with GSH values >2SD of group mean among the sample of patients. We repeated the regression analysis after excluding this outlier. The results of the regression predicting TTR indicated the two predictors (glutamate and GSH) explained 28% of the variance (R^2^ =.0.28, F(2,23)=4.13, p=0.031). Higher levels of GSH predicted a shorter time to response (β = -0.58, p=0.017) while glutamate was not a significant predictor (β = 0.09, p=0.68). The results of the regression predicting SOFAS at baseline indicated the two predictors explained 33% of the variance (R^2^ =.0.33, F(2,23)=5.08, p=0.016). Higher levels of glutamate predicted lower SOFAS scores (β = -0.63, p=0.007) while GSH was not a significant predictor (β = 0.13, p=0.53). Thus, our regression analyses were robust to the effect of outliers.

### Clinical relevance

**Prognostic relevance:** To provide clinically relevant information to a patient with FEP whose GSH values have been quantified as described in this study, we performed a median split of the patient group based on ACC GSH values. We then compared the low-GSH and high-GSH FEP groups on the clinically relevant variables of time to response; percentage symptom reduction and time to reach CGI score of 2 (much improved).

|  | Low GSH group (n=12) | High GSH group (n=13) |
| --- | --- | --- |
| Time to 50% symptom reduction (weeks) | 9.25(6.53) | 4.15(2.27) |
| Time to achieve CGI score 2 (weeks) | 8.41 (6.4) | 3.92 (2.6) |
| Total symptom reduction (%) | 29.3% (42%) | 67.4% (26%) |

Patients with FEP who have higher GSH levels will reach 50% symptom reduction at a rate that is 55% faster than those with lower levels when taking antipsychotics (absolute difference ~5.5 weeks); in terms of the symptom scores, they achieve 57% more reduction in symptom burden with 4 weeks of antipsychotic therapy.

**Therapeutic Relevance:** Conus et al (2018)^14^ have demonstrated that 6 months of NAC supplementation at a dose of 2700mg/day results in 28% increase in prefrontal glutathione levels in patients with first episode psychosis. They noted that nearly 70% of all patients receiving NAC show >10% increase in baseline GSH levels with NAC supplementation.

We extrapolated this information to estimate the degree of benefit we could expect in our clinical sample if the patients with low GSH levels receive NAC supplementation.

We first converted GSH levels to percentile rank values. Following this we repeated the regression analysis predicting time-to-response as reported in the manuscript, as well as the actual percentage reduction in PANSS-8. We noted that for every 1% increase in GSH levels, we could see 0.11 weeks of reduction in time to response and 0.67 points reduction in PANSS-8. In other words, if we can increase GSH levels by 10% in this sample, we can expect response (50% reduction in psychotic symptoms) to be achieved 7 days earlier than expected. In terms of actual symptom levels, a 7% extra reduction in baseline PANSS-8 burden can be expected for each 10% increase in GSH levels.

### The use of PANSS-8 to assess clinical outcome

Given the acute stage of recruitment, and our emphasis on obtaining clinical scores on the day of scanning, we did not obtain PANSS-30 measurements, and relied on PANSS-8 for estimating the clinical trajectory. In a head-to-head comparison of PANSS-8 and PANSS-30 in a sample of 270 subjects in an acute phase of schizophrenia, the correlation between PANSS-8 and PANSS-30 was noted to be excellent across various time points [r=0.85 (baseline), 0.88 (1 week), 0.89(2 weeks),0.90 (3 weeks), 0.90(4 weeks), 0.91 (6 weeks)]^1^. More importantly, when comparing the sensitivity to change for PANSS-8 and PANSS-30 using T scores, a remarkable similarity over time has been observed (See below an excerpt from Table 1 of Lin et al., 2018). Given this data, we estimate the likelihood of misclassification when using PANSS-8 instead of PANSS-30 in our sample is very low.

In terms of cross-sectional remission, we observed the rates of 42.31% (n=11 of 26) a 1-month, 50% (n=13 of 26) at 3-months and 60% (n= 15 of 25) at 6 months. Our observed rates of remission are comparable to Simonsen et al^2^., who reported 56.5% cross-sectional remission at 3-months in a comparable consecutive sample of FEP (n=301, PANSS P1, P3, P5, P6 or G9 <4), and 45.6% early response rate by Verma et al^3^., (n=1175, using PANSS-30, 40% improvement criteria), and 53.3% reported by Nordon et al^4^. (n=467, using CGI<4 and >30% improvement criteria). Our observation that remission rates increased from from 42.31% at 4 weeks to 60% at 24 weeks also concurs with numerous other observations suggesting that patients with FEP are still improving beyond 1-month duration^5–8^. Thus a longer period of evaluation is crucial in first-episode samples before we can consider treatment failure or non-remission.

### Antipsychotics, clinical outcome and MRS measures

To examine the effect of baseline exposure to antipsychotics, we included the total antipsychotic dose exposure at the time of scanning (in DDD dose year equivalents) as a covariate and tested the effect in our regression analyses. The results of the regression predicting time to response (TTR) indicated the 3 predictors (glutamate, GSH and DDD) explained 37% of the variance (R2 =.0.37, F(3,24)=4.11, p=0.019). Higher levels of GSH predicted a shorter time to response (β = -0.54, p=0.048) while glutamate (β = 0.06, p=0.81) and antipsychotic dose (β = 0.26, p=0.16) were not significant predictors. The results of the regression predicting SOFAS at baseline indicated the two predictors explained 36% of the variance (R2 =.0.36, F(3,24)=3.96, p=0.022). Higher levels of glutamate continued to predict lower SOFAS scores (β = -0.65, p=0.015) while GSH (β = 0.16, p=0.51) and antipsychotic dose (β = 0.19, p=0.30) were not significant predictors. Thus, the reported relationships between GSH and glutamate with TTR and SOFAS scores respectively, are not explained by variations in the dose of antipsychotic exposure.

To examine the effect of injectable vs. oral antipsychotics on clinical outcome , we studied the effects of using long-acting injectables (n=12) vs. oral antipsychotics (n=14) during the course of treatment. There were no differences in time to achieve 50% PANSS-8 reduction [TTR] (LAI users mean (SD) = 5.92(3.5); non-users mean (SD) = 7.23(6.7); p=0.55) or time to achieve CGI score of 2 (LAI users mean (SD) = 6.08(5.5); non-users mean (SD) = 6.07(5.2); p=0.99) levels, or the percentage reduction in PANSS-8 at 1-month follow-up (LAI users mean (SD) = 54.2%(27.4%); non-users mean (SD) = 45.2%(50.2%); p=0.60) between patients who were treated with LAIs by 1 month and those who continued with oral antipsychotics. It is important to note that LAIs were used only after at least 2 weeks of oral antipsychotic administration (as per the product monographs), explaining the lack of notable differences in the rates of response between the LAI and non-LAI users.

To examine the effect of partial agonists vs. D2-blockers on clinical outcome, we studied the effects of using partial agonists (aripiprazole oral or LAIs and brexpiprazole, n=7) vs. other D2-antagonists (n=19) during the course of treatment. There were no differences in time to achieve 50% PANSS-8 reduction [TTR] (partial agonist users mean (SD) = 3.71(2.1); non-users mean (SD) = 7.23(5.9); p=0.09) or time to achieve CGI score of 2 (partial agonist users mean (SD) = 3.71(1.6); non-users mean (SD) = 7.0(5.8); p=0.16) levels, or the percentage reduction in PANSS-8 at 1-month follow-up (partial agonist users mean (SD) = 74.5%(21.3%); non-users mean (SD) = 40.9%(40.1%); p=0.07) between patients who were treated with partial agonists and those who received D2 antagonists as antipsychotics. Given the small number of patients who received partial agonists (n=7), we urge caution in extrapolating these results outside of the context of this study. Nevertheless, these results indicate that speed of response (TTR) is not confounded by difference in the choice of antipsychotics in this sample.

### Substance use, clinical outcome and MRS measures

The determination of substance use disorder in the past year was based on self-report questionnaire for controls, and self-report, DSM-based clinical assessment as well as urine drug screening at the point of clinic entry (not on the day of scanning) in suspected cases for patients. None of our subjects satisfied substance use disorder criteria in the past year as per DSM-5.

In addition to this DSM based exclusion criteria, we screened all subjects for recreational drug use using a self-report substance use questionnaire comprised of AUDIT-C (6 months history) for alcohol use^6^, smoking index (in pack years = years of regular smoking X number of cigarettes per day/20), single item of Fagerström Test for Nicotine Dependence (time to first cigarette in the morning^7^), and 6-item Cannabis Abuse Screening Test (CAST^8^) and the 10-item Drug Abuse Screening Test (DAST-10^9^) for substances other than cannabis, alcohol and nicotine.

Among the 18 patients who endorsed smoking nicotine regularly, we found no correlation between smoking index and glutamate (Spearman’s rho =-0.01, p=0.9) or glutathione (rho = -0.13, p=0.64). We did not have sufficient data to explore the effect of alcohol use (only 5 patients and none of the controls had AUDIT-C score >4).

To address the possible effect of cannabis on MRS data, we first compared 18 patients who endorsed recreational cannabis use and the 9 non-users. There were no differences in glutamate (cannabis users mean (SD) = 9.05(2.16); non-users mean (SD) = 7.30(1.12); p=0.06) or glutathione (cannabis users mean (SD) = 1.78(0.44); non-users mean (SD) = 1.64(0.25); p=0.49) levels, or time to response (cannabis users mean (SD) = 5.38(3.59); non-users mean (SD) = 9.19(7.62); =0.12) between patients who were self-identified cannabis-users and non-users.

Among the 18 patients who endorsed recreational use of cannabis, we found no correlation between total CAST scores and glutamate (r=0.36, p=0.14) or glutathione (r=0.17, p=0.51). We also undertook a median split of the cannabis using patient sample based on CAST scores (heavy users >14 or low-level users <14, n=9 in each group) and noted no differences in time to response (TTR; heavy users mean (SD) = 5.1(3.9); low-level users mean (SD) = 9.5(7.2);, p=0.12), glutamate (heavy users mean (SD) = 9.4(2.4); low-level users mean (SD) = 8.3(2.1); p=0.31) or glutathione (heavy users mean (SD) = 1.92(0.5); low-level users mean (SD) = 1.6(0.3); p=0.12) levels. But given the small numbers of recreational cannabis users, we cannot conclusively rule out the possibility that cannabis use has an impact on MRS metabolites of interest.

### References

1.	Lin, C.-H. *et al.* Early improvement in PANSS-30, PANSS-8, and PANSS-6 scores predicts ultimate response and remission during acute treatment of schizophrenia. *Acta Psychiatr. Scand.* **137**, 98–108 (2018).

2.	Simonsen, E. *et al.* Early identification of non-remission in first-episode psychosis in a two-year outcome study. *Acta Psychiatr. Scand.* **122**, 375–383 (2010).

3.	Verma, S., Subramaniam, M., Abdin, E., Poon, L. Y. & Chong, S. A. Symptomatic and functional remission in patients with first-episode psychosis. *Acta Psychiatr. Scand.* **126**, 282–289 (2012).

4.	Nordon, C. *et al.* Trajectories of antipsychotic response in drug-naive schizophrenia patients: results from the 6-month ESPASS follow-up study. *Acta Psychiatr. Scand.* **129**, 116–125 (2014).

5.	Schennach-Wolff, R. *et al.* Predictors of response and remission in the acute treatment of first-episode schizophrenia patients--is it all about early response? *Eur. Neuropsychopharmacol. J. Eur. Coll. Neuropsychopharmacol.* **21**, 370–378 (2011).

6.	Derks, E. M. *et al.* Antipsychotic drug treatment in first-episode psychosis: should patients be switched to a different antipsychotic drug after 2, 4, or 6 weeks of nonresponse? *J. Clin. Psychopharmacol.* **30**, 176–180 (2010).

7.	Emsley, R. *et al.* Remission in first-episode psychosis: predictor variables and symptom improvement patterns. *J. Clin. Psychiatry* **67**, 1707–1712 (2006).

8.	Stentebjerg-Olesen, M. *et al.* Early nonresponse determined by the clinical global impressions scale predicts poorer outcomes in youth with schizophrenia spectrum disorders naturalistically treated with second-generation antipsychotics. *J. Child Adolesc. Psychopharmacol.* **23**, 665–675 (2013).
